# The function of human PIF1 in G quadruplex formation and replication stress response at ALT telomeres

**DOI:** 10.64898/2026.08.31.748286

**Authors:** Ashwin Ragupathi, Rongwei Zhao, Shakir Najit, Anthony Fernandez, Heba Abid, Jiho Mo, Samantha Juang, Christopher Tong, Derin I. Yetil, Liam Buchanan, Connor McDowell, Lindsay Mosca, Bronwyn Kelly, Rebecca Kelly, Zakir Hossain, Subhabrata Chaudhury, Jaewon Min, Huaiying Zhang, Ming Xiao, Binghui Shen, Dong Zhang

**Affiliations:** Department of Biomedical Sciences, College of Osteopathic Medicine, New York Institute of Technology, Old Westbury, NY 11568, USA; Center for Cancer Research, New York Institute of Technology, Old Westbury, NY 11568, USA; Department of Biology, Carnegie Mellon University, Pittsburgh, PA 15213, USA; Department of Cancer Genetics and Epigenetics, Beckman Research Institute, City of Hope, Duarte, CA 91007, USA; School of Biomedical Engineering, Science and Health System, Drexel University, Philadelphia, PA 19104, USA; Institute of Cancer Genetics, Columbia University Vagelos College of Physicians and Surgeons, NY, NY 10032, USA; Department of Biological and Chemical Sciences, New York Institute of Technology, Old Westbury, NY 11568, USA; Center for Genomic Sciences and Center for Advanced Microbial Processing, Institute of Molecular Medicine and Infectious Disease, Drexel University College of Medicine, Philadelphia, PA, 19102, USA

## Abstract

Cancers maintain their telomeres through two telomere maintenance mechanisms: 85-90% of cancers rely on telomerase (TEL+), while 10-15% of cancers adopt the Alternative Lengthening of Telomeres (ALT) pathway. The Break-Induced Replication (BIR) pathway plays a critical role in maintaining telomere length in the ALT+ cells. In both yeast and human, PIF1, a 5’ to 3’ helicase, is required for the robust activity of BIR. However, the extent of human PIF1 (hPIF1) involvement in the ALT pathway remains unknown. Here we showed that hPIF1 can be recruited to damaged telomeres in ALT+ cells. In addition, we demonstrated that inhibition of hPIF1 induced DNA damage and G quadruplex (G4) accumulation at ALT telomeres, leading to a moderate reduction of the mean telomere length. Most interestingly, we demonstrated that inhibition of hPIF1 also attenuates checkpoint activation, BLM recruitment, single-stranded DNA (ssDNA) formation, DNA damage, and G4s at telomeres in the FANCM deficient ALT+ cells. Finally, we showed that inactivation of hPIF1 affects the viability of both ALT+ and TEL+ cancers, suggesting that hPIF1 is a potential drug target for cancer therapy.

## Introduction

Human telomeres, the repetitive nucleoprotein structures located at the ends of every linear chromosomes with the canonical sequence of (TTAGGG)_n_, are crucial for maintaining the integrity of our genome. To achieve unlimited proliferation, most tumor cells adopt a telomere maintenance mechanism. While 85-90% of human cancers reactivate telomerase (TEL), 10-15% of them rely on the Alternative Lengthening of Telomeres (ALT) pathway(Bryan et al., 1997; Kim et al., 1994; MacKenzie et al., 2021; Shay and Bacchetti, 1997). Due to their unique structures and sequences, telomeres are considered the difficult-to-replicate genomic regions and are prone to replication stress and spontaneous DNA damage, particularly in ALT-positive (ALT+) cells(Cesare et al., 2009; Doksani, 2019; Douglas, 2024; Ragupathi et al., 2023). For example, the G-rich strand of telomeres can form G-quadruplexes (G4s)(Drosopoulos et al., 2015). The telomeric long noncoding RNA, TERRA, can be transcribed from almost all the chromosome arms(Hsieh et al., 2025; Rodrigues et al., 2024). Pairing of TERRA with the C-rich strand, either via cis- or trans-, can lead to the formation of TERRA R-loops, which are elevated in ALT+ cells(Arora et al., 2014; Azzalin et al., 2007). Failure to unfold G4s and remove TERRA R-loops will impede replisome progression, ultimately leading to replication fork stalling/collapse and spontaneous DNA damage at telomeres.

In the absence of telomerase, ALT+ cells hijack a cohort of homology-dependent repair (HDR) factors to maintain their telomere length and avoid cell death triggered by severe telomere attrition(O’Sullivan and Greenberg, 2025). Moreover, because of their unique chromatin properties and the tendency to experience heightened spontaneous DNA damage, ALT telomeres have acquired additional strategies to suppress replication stress. For example, we and others have shown that the Fanconi Anemia complementation group M protein (FANCM) is needed to actively remove TERRA R-loops at ALT telomeres, but not at TEL telomeres, and facilitates telomere replication(Lu et al., 2019; Pan et al., 2019; Pan et al., 2017; Silva et al., 2019). In addition to the absence of telomerase activity, a constellation of additional molecular and cellular biomarkers helps to define the ALT-positivity(MacKenzie et al., 2021). The biomarkers that help to distinguish ALT+ cell from TEL+ cells include highly heterogeneous telomere lengths, increased telomere sister chromatid exchange (tSCEs), the presence of unique telomeric sequence variants (TSV), increased telomere dysfunction-induced foci (TIFs), the formation of ALT-associated PML bodies (APBs), and a drastic increase of extrachromosomal telomeric repeats (ECTRs), most notably C-circles(MacKenzie et al., 2021). Recently, we demonstrated that the Single-Molecule Telomere Assay via Optical Mapping (SMTA-OM) can be also used to differentiate the ALT+ from the TEL+ cells while the Reddel’s and MacKenzie’s groups derived ALT- and TEL-predictors from the proteomic and transcriptomic analysis(Raseley et al., 2023; Wu et al., 2025b).

Break-Induced Replication (BIR) is a specialized HDR pathway and plays a critical role in break-induced telomere synthesis (BITS) and ALT pathway(Dilley et al., 2016; Lu et al., 2019; Pan et al., 2019). BIR was initially discovered and well characterized in budding yeast, *Saccharomyces cerevisiae*, and was primarily used to repair a single-ended double-stranded DNA break (seDSB). Haber and colleagues demonstrated that Pol32 (PolD3 in human), a subunit of DNA polymerase delta (Polδ) and a nonessential gene in budding yeast, is required for BIR(Lydeard et al., 2007). Subsequently, two groups showed that in budding yeast, BIR also requires PIF1, a 5’ to 3’ DNA helicase, which facilitates Polδ-dependent DNA synthesis through a bubble migration mechanism(Chung et al., 2010; Saini et al., 2013; Wilson et al., 2013). Recently, using EGFP-based BIR reporters, Wu and colleagues showed that human PIF1 (hPIF1) also plays a critical role in BIR(Li et al., 2021). However, involvement of hPIF1 in the ALT pathway remains largely unknown.

Here, we investigated the role of hPIF1 in the ALT pathway. We first demonstrated that hPIF1 can be recruited to damaged ALT telomeres. We then showed that inhibition of hPIF1 induced G4 accumulation, replication stress, and DNA damage at ALT telomeres. Furthermore, we showed that hPIF1 facilitates the checkpoint activation and suppresses the formation of single-stranded DNA (ssDNA) and DNA damage in FANCM deficient ALT+ cells. Finally, we showed that inactivation of hPIF1 affects the viability of both ALT+ and TEL+ cancer cells, suggesting that hPIF1 is a potential drug target for cancer therapy.

## Materials and Methods

### Cell lines, tissue culture, and chemicals

A549, H460, HeLa, U2OS and Saos2 were purchased from ATCC. GFP-hPIF1 was generously provided by Dr. Stephen Jackson. All cells are grown in DMEM (Corning) supplemented with 10% fetal bovine serum (Bio-techne), penicillin (Corning), and streptomycin (Corning), at 37 °C in a humidified incubator with 5% CO_2._ PIF1i-C1 was purchased from AmBeed and dissolved in DMSO. The purity of PIF1i-C1 was validated in-house through HPLC, mass spectrometry, 1H_NMR, and 13C-NMR (Fig S2).

### TRF1-FokI induction

GFP-hPIF1/U2OS cells were first transfected with mCherry-ER-DD-TRF1–FokI (WT) or mCherry-ER-DD-TRF1–FokI (D450A). Shield-1 (Cheminpharma LLC) and 4-hydroxytamoxifen (Sigma-Aldrich) at 1 μM were then added for 2 hours to allow TRF1 to enter the nucleus after first-round imaging, as previously described (Cho et al., 2014).

### Cell imaging and image processing

Imaging acquisition was performed as previously described(Xu et al., 2024). For live imaging, cells were seeded on 22 × 22-mm glass coverslips coated with poly-D-lysine (P1024; Sigma-Aldrich). When ready for imaging, coverslips were mounted in magnetic chambers (Chamlide CM-S22-1; LCI) with cells maintained in a normal medium supplemented with 10% FBS and 1% penicillin/streptomycin at 37°C on a heated stage in an environmental chamber (TOKAI HIT Co., Ltd.). Images were acquired with a microscope (ECLIPSE Ti2) with a 100 × 1.4 NA objective, a 16 XY Piezo-Z stage (Nikon Instruments, Inc.), a spinning disk (Yokogawa), an electron multiplier charge-coupled device camera (IXON-L-897), and a laser merge module that was equipped with 488, 561, 594, and 630 nm lasers controlled by NIS-Elements Advanced Research. For both the fixed cells and live imaging, images were taken with 0.5 μm spacing between Z slices, for a total of 8 μm. For movies, images were taken at 5-min intervals for up to 3 h. All error bars represent means ± SEM. Statistical analyses were performed using Prism 10.0 (GraphPad software). Two-tailed unpaired *t* tests have been used for all tests. Statistical significance: P > 0.05; *P < 0.05; **P < 0.01; ***P < 0.001.

### siRNA transfection

Cells were transfected with 100 nM of siRNA (Supplemental Table 2) and RNAiMax (13778075; Thermo Fisher Scientific) diluted in OptiMEM (31-985-070; Life Technologies). The transfection medium was replaced with culture media 6 hours later, and the transfection was repeated on Day 2. Cells were collected and imaged at 48 hours after the second-round transfection. The sequence and order information for all the siRNA are listed in the Supplemental Table-3.

### Immunoblotting

Cells are collected and lysed directly in a sample buffer to achieve a concentration of 10^4^ cells/μl, followed by sonication for 30 seconds. Equal volumes of lysate were loaded for immunoblotting (Supplemental Table 3).

### Immunofluorescent staining

Cells were seeded onto coverslips. Cells were either then fixed with 3% paraformaldehyde containing 2% sucrose for 10 min, followed by treatment with Triton X-100 solution on ice for 5 min, or treated with Triton X-100 solution on ice for 5 min then fixed with 3% paraformaldehyde containing 2% sucrose for 10 min (pre-extraction). Slides were then blocked with 1% Gelatin in PBS. Cells are stained with primary antibodies and the respective Alexa-488 (Invitrogen) and Alexa-546 (Invitrogen) conjugated secondary antibodies. All the antibodies used for immunofluorescent staining are listed in Supplemental Table 3. SlowFade Gold DAPI (S36938) was used to stain the DNA. Images were collected by using an Olympus upright Fluorescent Microscope images were processed using Adobe Photoshop.

### C-circle assay

Cells were transfected twice with different siRNA. 48 hours later, genomic DNA from 3-5 x 10^5^ cells was extracted using QIAamp DNA Blood Mini Kit (Qiagen, 51106). 4 μg genomic DNA were digested with Alu I (NEB, R0137L) and Mbo I (NEB, R0147L) at 37 °C for at least 2 hours and then purified using Qiagen PCR Purification Kit (Qiagen, 28106). 40 ng of Alu I and Mbo I digested DNA were used for the Phi29 DNA polymerase reaction (30 °C for 8 hours and then 65 °C for 20 min). All the PCR reaction mixtures were loaded onto the Amersham Hybond-N+ membrane using the Bio-Rad Bio-Dot Microfiltration apparatus. The membrane was cross-linked using the UV Stratalinker at energy 1200. The membrane was then processed, probed, and developed using the TeloTAGGG Telomere Length Assay kit (Roche, 12-209-136-001). Images were taken with an Amersham Imager 600 and quantified using the NIH ImageJ.

### Strand-Specific Southern for Single-stranded Extrachromosomal Telomeres (4SET) assay

4SET was done as described previously(Lee et al., 2024).

### Single-molecule telomere assay via optical mapping (SMTA-OM)

Cells were transfected with different siRNAs, and stored in 10% DMSO in liquid nitrogen until DNA extraction and Purification.

#### DNA extraction and purification

Cells were first embedded in gel plugs with approximately 1 million cells per gel plug (BioRad no. 170-3592). The high molecular weight genomic DNA was extracted and purified using the Bionano Genomics SP kit.

#### DNA Labeling

Two motif-mapping approaches were used for the first part of the DNA labeling procedure. (1) The Nt.BspQI protocol, which was used for U2OS cells: The DNA sample was labeled as previously described (McCaffrey et al., 2017). (2) The DLE-1 protocol, which was used only for the Saos2 cells: Following the manufacturer’s instructions, a DLS labeling kit (Bionano Genomics) labeled 750 ng of genomic DNA. A labeling mix comprised Direct Labeling Enzyme 1 (DLE-1), 1X DLS reaction buffer, and DL green fluorophore-labeled nucleotide mix. The labeling mix was added to the genomic DNA, gently mixed via a wide bore pipette, and then incubated at 37°C for 2 hours. Following the incubation, membrane dialysis was used to remove the unwanted fluorescent dyes, proteins, and salts. Membrane dialysis was done at room temperature for approximately 2 hours in the dark to protect it from the light. Dialyzed DNA was then recovered using a 100 nm hydrophilic membrane (EMD Millipore, VCWP04700) and quantified with a Qubit device.

The second part of the DNA labeling procedure focuses on labeling the telomeres. Guide RNA (gRNA) consisting of 0.5 μM tracrRNA (IDT) and 50 μM crRNA was gently mixed via pipetting up and down and annealed on ice for 30 minutes. 25 pmol gRNA was incubated with 200 ng Cas9D10A nicking enzyme and 1X NE Buffer 3.1 at 37°C for 15 minutes. Next, 300 ng of the previously DLE-1 labeled genomic DNA was added to the Cas9D10A mixture for the one-hour nicking reaction at 37°C. After nicking, 200 nM red fluorophore-labeled nucleotides (ATTO647-dUTP, dATP, dGTP, dCTP) were incorporated into telomeres by 5U Taq DNA polymerase at 72°C for one hour in 1X Thermopol buffer (New England Biolabs). The nick-labeled sample was treated with proteinase K (QIAGEN) at 50°C for 30 min. Finally, to prepare the DNA for nanochannels, a staining mix of flow buffer, DTT, and YOYO-1 (a blue-colored dye that labels the backbone of DNA, Bionano Genomics DLS kit) was prepared according to the manufacturer’s instruction before adding it to the sample. The sample combined with the staining mix was incubated at room temperature overnight.

#### Imaging with Saphyr

A Bionano Saphyr G1.2 chip was loaded with the labeled DNA sample and imaged using a “dual-labeled sample” scheme incorporated in the Saphyr software. Images were taken in the following order: first, the red fluorescent labeling with the 637 nm laser, then the green fluorescent labeling with the 532 nm laser, and finally, the blue fluorescent DNA backbone YOYO-1 staining with the 473 nm laser.

#### Genome Assembly

*De novo* genome assembly was done using the software developed by Bionano Genomics. Consensus maps were generated by de novo assembling individual DNA molecules followed by alignment to the hg38 human reference genome.

#### Telomere analysis

CMAP, XMAP, and BNX files were generated after *de novo* genome assembly. The CMAP files contain both the red and the green label information. The XMAP files contain only the information on the green labels. Molecules matching the expected green labeling patterns were extracted from the BNX and CMAP files. Raw molecule images were extracted using in-house software based on their locations in the nanochannels. The red telomere labels were easily distinguishable from the green labels in the subtelomeric region. Telomeres were then analyzed and measured using the ImageJ software. The ferret diameter tool was used to capture the length of telomeres in pixels and intensities before converting all the measurements to kilobase (kb) following the previously established procedures(Abid et al., 2020; McCaffrey et al., 2017).

### G4 staining in cell culture, immunostaining, FISH, and foci quantification

Cells were fixed using 4% PFA in PBS for 15min at RT, followed by 0.5% Triton X-100 (15min, RT), and blocking in 5% BSA + 0.1% Tween-20 and Image iT-FX Signal Enhancer. The anti-G4 primary antibody (Sigma, MABE1126, 1:500) was diluted 1:500 in 4% BSA and 0.1% Tween-20 overnight at 4 °C, washed with 0.1% Tween-20, and stained with 1:1000 GAM Alexa Fluor 568 (Thermo, A11031) for 1 h at RT. For imaging experiments where FISH was used, cells were dehydrated using an ethanol series dehydration and labeled with a TelG-Alexa 488 telomere probe (PNA Bio) at 37 °C for 2 h. After rinsing, cells were immunostained for G4 and the appropriate protein. Cells were finally counterstained with DAPI and mounted in ProLong Gold (Thermo). Imaging was performed using the Zeiss LSM 900 with Airyscan 2. Briefly, we zoomed into individual nuclei and scanned with a resolution that a theoretical point spread function would be sampled at least 3×. After acquisition, images were processed using Zeiss’s Airyscan Joint Deconvolution algorithm at 10 iterations to ensure maximum resolution. Telomeric G4 were measured first by finding telomere spots. ROIs of consistent size were positioned at each identified telomere spot, and G4 intensity in the G4 channel was measured at each telomere position. ROI sizes were roughly 250 nm x 250 nm. Statistical significance was established using Dunnett’s T3 multiple comparisons test. To quantify foci, we used the particle finder algorithm on ImageJ (1.54 f). We used a consistent threshold across all imaging conditions and measured the area of foci divided by the area of the nucleus over at least 40 cells and at least 3 biological replicates per condition.

### Crystal violet assay

4000 cells/per well are seeded in 12 well plates. After 7-14 days of growth, cells are fixed using a methanol/acetic acid solution and stained with 1% crystal violet solution.

## Results

### hPIF1 can be actively recruited to damaged telomeres

Because of the critical function of PIF1 in BIR (Chung et al., 2010; Li et al., 2021; Saini et al., 2013; Wilson et al., 2013), we hypothesized that hPIF1 is also involved in the ALT pathway. To test this, we first investigated telomere localization of hPIF1 using a U2OS cell line that stably expresses a GFP-tagged hPIF1 (GFP-hPIF1), which localizes exclusively in the nucleus (Rodriguez et al., 2012). U2OS is a commonly used cellular model for investigating the ALT pathway. In non-stressed U2OS cells, there is no pronounced localization of GFP-hPIF1 to telomeres (data not shown). We then examined whether hPIF1 can be detected at damaged telomeres. For this, we utilized the TRF1-FokI system developed by Greenberg and colleagues (Cho et al., 2014; Dilley et al., 2016). In this system, the restriction enzyme, FokI, which recognizes the consensus sequence, 5’-GGATG-3’, and cuts after the ninth nucleotide on the sense strand and after the thirteenth nucleotide on the antisense strand, is fused with TRF1, an important subunit of the Shelterin Complex(de Lange, 2018). Additionally, TRF1-FokI is also fused with a destabilization domain (DD), a modified estrogen receptor (ER) domain, and a red fluorescent protein, mCherry. The presence of Shield-1 prevents DD-mediated proteasome degradation. When 4-hydroxytamoxifen (4-OHT) is added to the culture medium, TRF1-FokI is translocated from cytoplasm to nucleus, binds most telomeres, and cuts them. Simultaneously, the damaged telomeres can be visualized through mCherry. An enzyme-dead FokI (FokI-D450A) is used as the negative control for the TRF1-FokI system because though FokI-D450A can be recruited to telomeres, it cannot cut them. Live cells were monitored using a time-lapse fluorescent microscope. As seen in Fig 1 and Video 1 and Vide 2, translocation of the wildtype TRF1-FokI (TRF1-FokI-WT) to telomeres, but not TRF1-FokI-D450A, induced a robust recruitment of GFP-hPIF1 to telomeres, indicating that hPIF1 can be actively recruited to the damaged telomeres.

**Figure 1.**
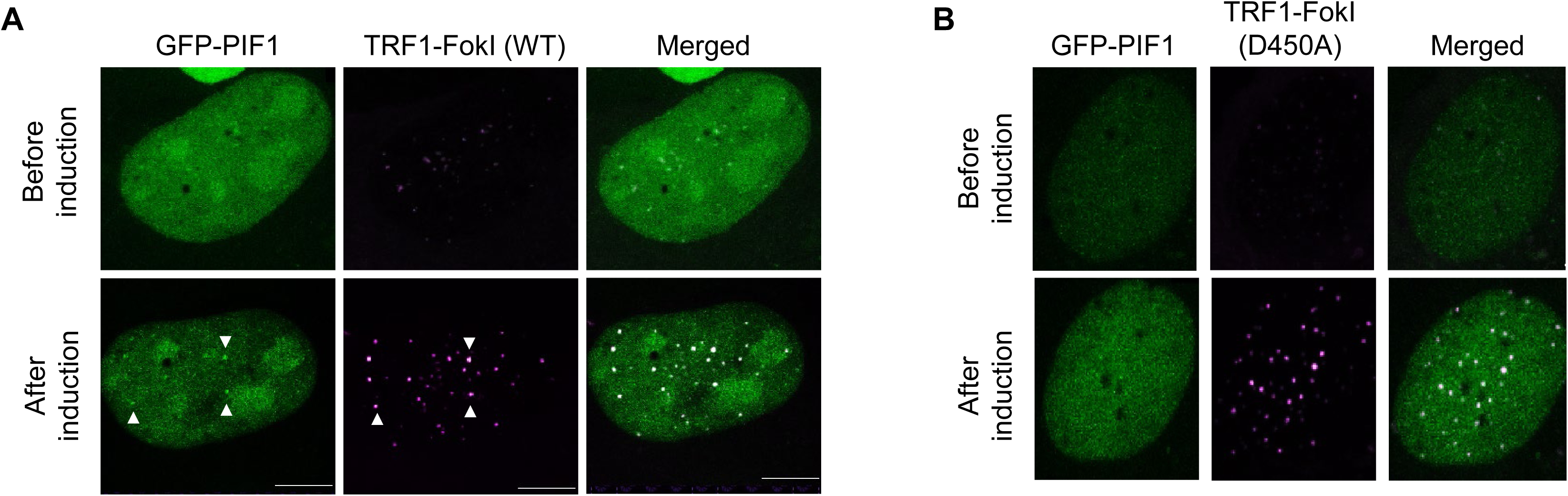
hPIF1 can be recruited to damaged ALT telomeres. Representative images of GFP-hPIF1 U2OS cells expressing either mCherry-TRF1-FokI-wildtype (WT)(A), or mCherry-TRF1-FokI-D450A (B).

### Inhibition of hPIF1 induced telomere damage and accumulation of C-rich ssDNA and C-circles in ALT+ cells

To investigate the role of hPIF1 in ALT pathway, we first inhibited the helicase activity of human PIF1 using a newly identified PIF1 inhibitor (Compound 1, or PIF1i), which was analyzed in-house using HPLC, mass spectrometry, and NMR and showed a purity close to 99% (Fig S1)(Wever et al., 2024). 72 hours after treating U2OS cells with 300 μM of PIF1i, we co-stained the cells with antibodies recognizing γH2AX, together with an antibody recognizing TRF2, another subunit of the Shelterin complex to mark the telomeres(de Lange, 2018). Interestingly, we observed a two-fold increase of γH2AX/TRF2 foci in the PIF1i treated cells (Figs 2A and 2B), indicating that inhibition of the helicase activity of hPIF1 induces DNA damage at ALT telomeres. Next, we depleted hPIF1 using two different siRNA in U2OS cells (Fig 2C). Transient depletion of hPIF1 led to a slight increase of G2 cells two days after the first siRNA transfection (Fig S2). We then imaged and quantified DNA damage at telomeres by co-staining cells with antibodies recognizing γH2AX together with the antibody recognizing TRF2. Consistent with the results from PIF1i treated U2OS cells, we also observed a pronounced increase of γH2AX/TRF2 foci in both U2OS (Fig 2D) and Saos2 (Fig 2E), another ALT+ cell line, suggesting that depletion of hPIF1 in ALT cells induced DNA damage at their telomeres. Finally, we monitored the C-rich ssDNA formation using the recently developed 4SET assay(Lee et al., 2024). Interestingly, we observed a reduction of the C-rich ssDNA in both siPIF1-1 transfected and PIF1i treated U2OS cells (Figs 3A to 3D). Consistently, we also observed a reduction of C-circle formation in those cells (Figs 3E to 3G). These data indicate that hPIF1 promotes the formation of ECTRs in ALT+ cells.

**Figure 2.**
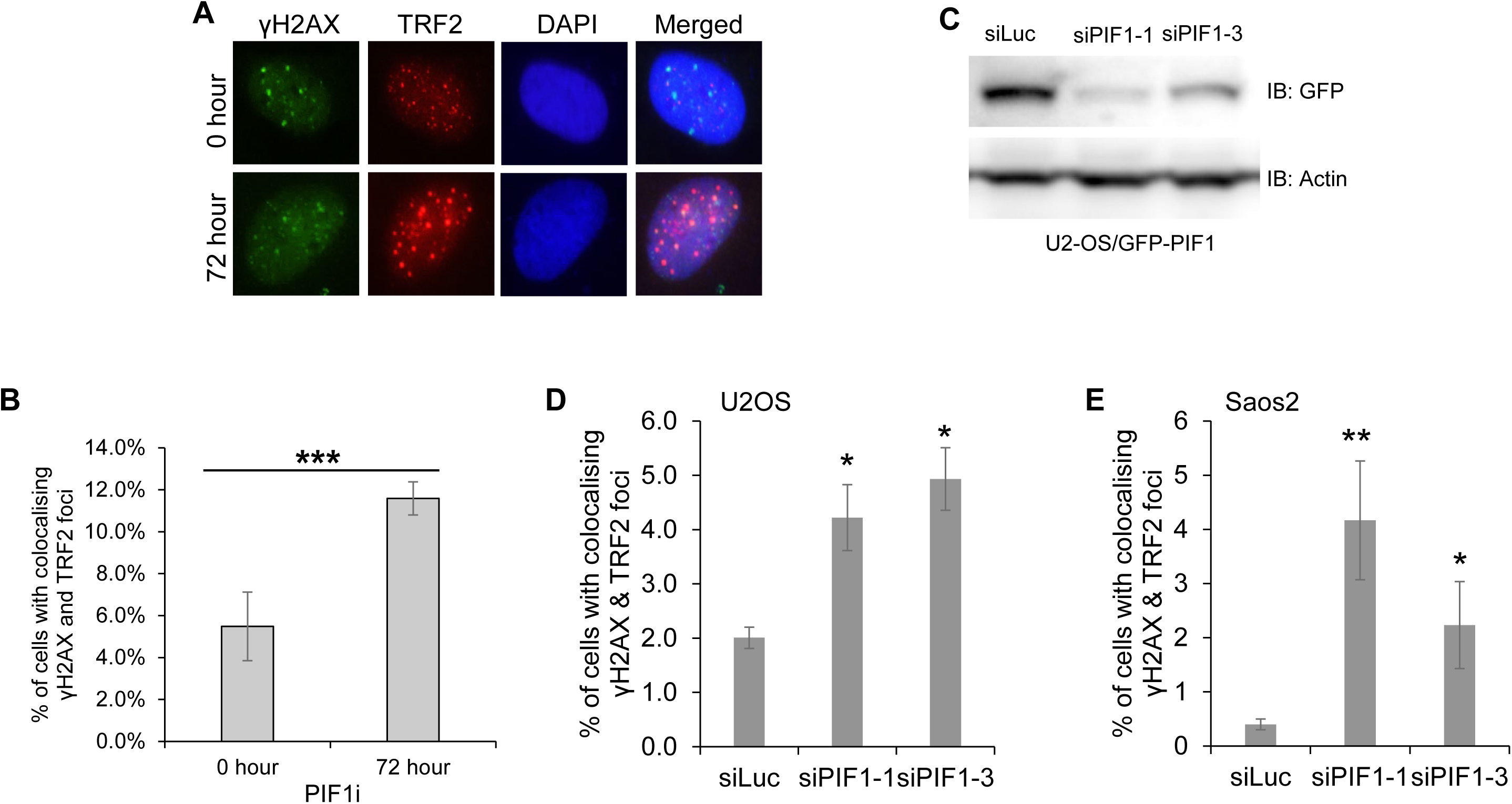
Inactivation of PIF1 induces DNA damage at ALT telomeres. (A and B) U2OS cells were treated either with DMSO (0 hour), or with 300 μM PIF1i for 72 hours, and then stained with antibodies recognizing TRF2 and γH2AX. U2OS cells (C and D) or Saos2 cells (E) were first transfected with siRNA, and then stained with antibodies recognizing TRF2 and γH2AX. All nuclei were stained with DAPI. All error bars are standard deviations obtained from three different experiments. Standard two-sided t test: \**p*<0.05, \*\**p*<0.01, \*\*\**p*<0.001.

**Figure 3.**
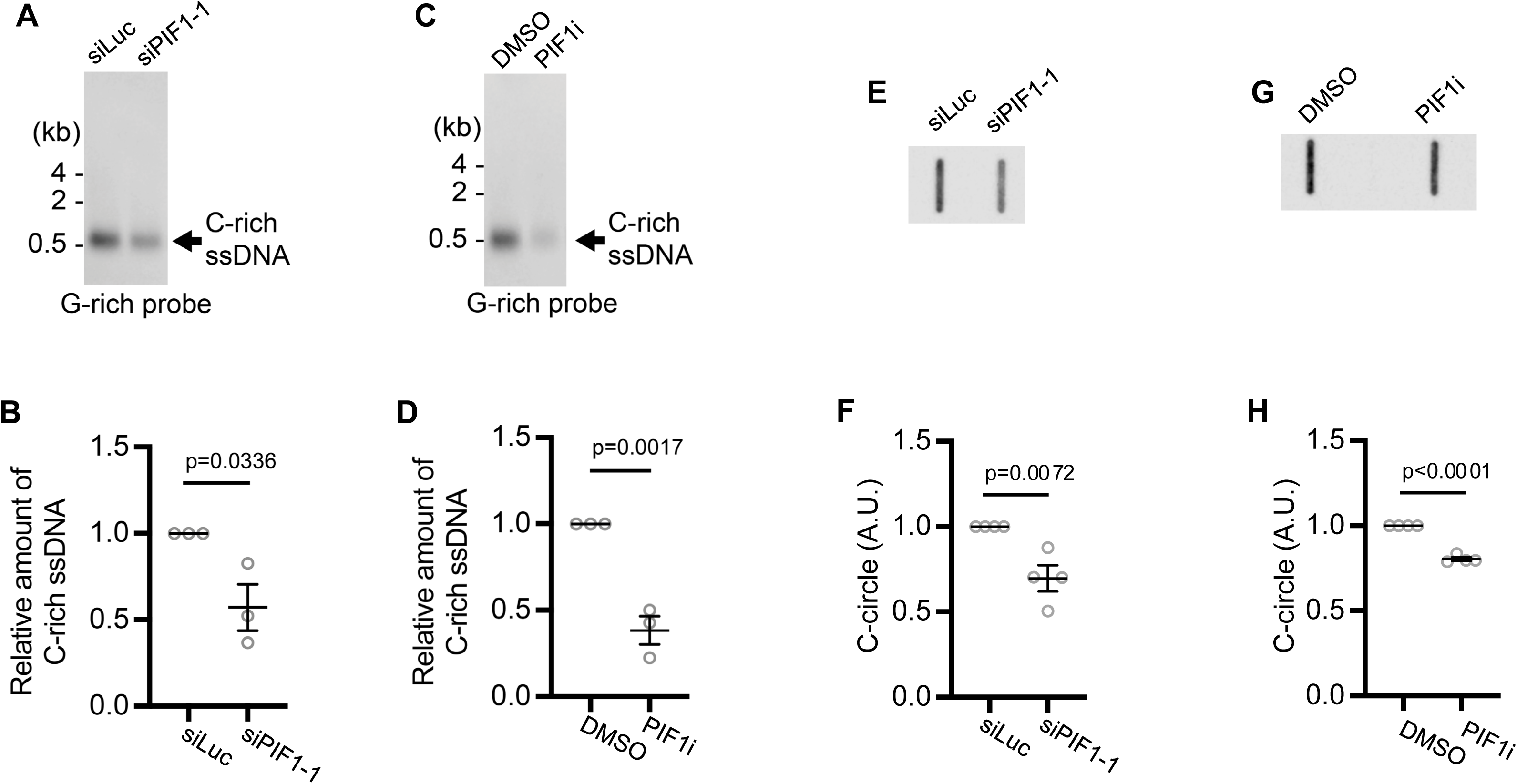
Inactivation of hPIF1 decreases telomeric C-rich ssDNA and C-circle formation in ALT cells. U2OS cells were transfected either with a control siRNA (siLuc), or a siRNA targeting hPIF1 (siPIF1-1), or were treated either with DMSO or with 300 μM PIF1i for 72 hours. DNA was extracted and used for 4SET assay (A to D) and C-circle assay (E to H). All error bars are standard deviations obtained from three or four different experiments. Standard two-sided t test: \**p*<0.05, \*\**p*<0.01, \*\*\**p*<0.001.

Collectively, our data indicate that hPIF1 plays a critical role in suppressing DNA damage at ALT telomeres while promoting the production of various ECTRs.

### Depletion of hPIF1 reduces mean end telomere length and the length of end telomeres at many chromosome arms while increasing chromosome fusions

Next, we measured telomere length in hPIF1 deficient U2OS cells using the SMTA-OM technology(McCaffrey et al., 2017; Raseley et al., 2023; Singh et al., 2024). Briefly, carefully extracted large genomic DNA molecules, averaging around 300 kb, are either nicked by a nickase Nt. BspQI (5’-GCTCTTCN-3’) and then labeled, or labeled directly by a direct-labeling enzyme (DLE-1, 5’-CTTAAG-3’, Bionano Genomics). The motif sequences identified by Nt. BspQI or DLE-1 throughout the genome are labeled with a green fluorophore. For labeling telomeres, DNA was first nicked by CRISPR/Cas9D10A and guided by a telomere guide RNA (gTelo) and then also labeled with a green fluorophore (McCaffrey et al., 2016; McCaffrey et al., 2017). All DNA molecules were co-stained with YOYO-1 (blue). The two-color (green and blue) labeled DNA molecules were then linearized in the NanoChannel Arrays and imaged. The patterns of spaced green labels from Nt. BspQI or DLE-1 were used to identify a specific chromosome arm by matching them to the predicted patterns of the reference genome, Hg38. A more intense and uniform green signal was identified as a telomere (Fig 4A). As reported previously, SMTA-OM enables not only the identification and measurement of telomere length in a chromosome arm-specific manner, but also the detection and quantification of additional features at the chromosome ends, including telomere free ends (TFE) and chromosome fusions, such as Fusion/ITS+ (ITS+)(Fig 4A)(Abid et al., 2020; Raseley et al., 2023). ITS+ manifests as a DNA molecule that on one side of the telomere signal, the DNA fragment is long enough to be assigned to a specific chromosome arm based on the pattern of its green labels. However, on the other side of the telomere signal, because the DNA fragment is too short, it cannot be confidently assigned to a particular chromosome arm based on the pattern of its green labels.

**Figure 4.**
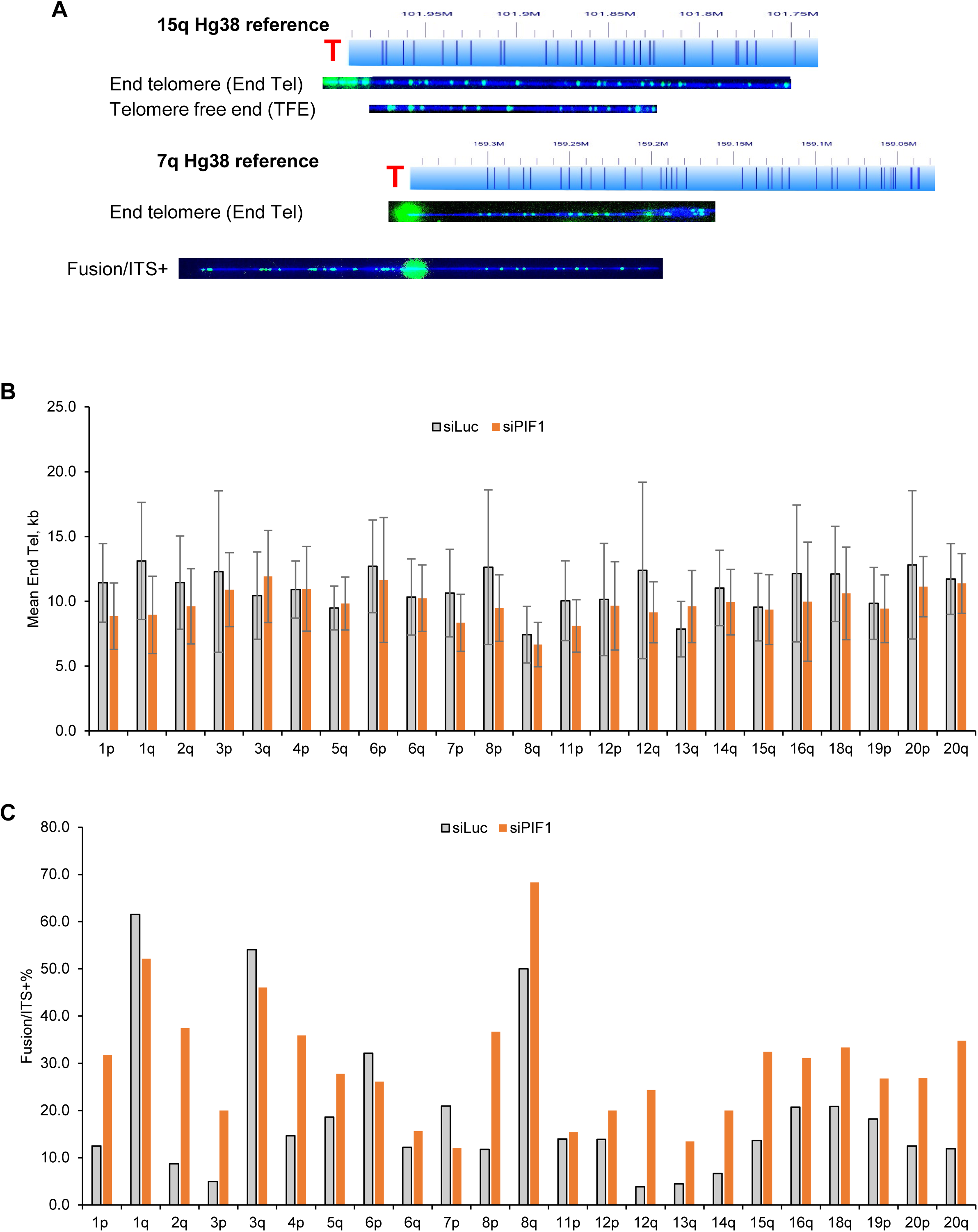
Inactivation of hPIF1 moderately decreases the mean length of telomeres. U2OS cells were first transfected with siLuc, or siPIF1-1. Genomic DNA were extracted from these cells and used for SMTA-OM assay. (**A**) Representative images of DNA molecules assigned to chromosome 7q and 15q. (**B**) The mean length of End telomere (End Tel) of different chromosome arms. (**C**) The percentage of Fusion/ITS+ at different chromosome arms.

The advantages of using SMTA-OM to define the features of telomeres and chromosome ends include: (1) there is no need to enrich telomeres, therefore can avoid the bias toward certain configuration of telomeres; (2) The detected and imaged DNA fibers can be quite long, ranging from 150 kb to 300 kb, which make it possible to confidently assign an end telomere (End Tel) to a particular chromosome arm; (3) chromosome fusions, including Fusion/ITS+ and Fusion/ITS-, can be easily detected and identified. We imaged and analyzed more than one thousand DNA molecules from siLuc or siPIF1-1 transfected U2OS cells. The mean End Tel length in the siPIF1-1 transfected cells showed a 12% reduction compared to the siLuc transfected cells (Table 1 and Supplemental Tables 1 and 2). Interestingly, the Fusion/ITS+ in the siPIF1-1 transfected cells showed a dramatic 71% increase compared to the siLuc transfected cells. Because there are only two telomere-free end (TFE) DNA molecules that are identified in the siPIF1-1 transfected cells out of the 1011 molecules imaged and analyzed, we are unable to calculate the p-Value for the TFEs. Nonetheless, there is a ten-fold reduction of the TFEs in the siPIF1-1 transfected cells. We speculate that though PIF1 deficiency may produce more TFEs, most of the TFEs are quickly converted to chromosome fusions with other damaged telomeres, forming Fusion/ITS+. When we examined telomere length at the level of individual chromosome arms, we observed that 13 arms showed reduction in the mean telomere length (1p, 1q, 2q, 3p, 7p, 8p, 8q, 11p, 12q, 14q, 16q, 18q, 20p), while six arms showed no change (4p, 5q, 6q, 12p, 15q, 20q) and 2 arms showed an increase (3q, 13q) (Fig 4B). Similarly, we observed that 19 arms (1p, 2q, 3p, 4p, 5q, 6q, 8p, 8q, 11p, 12p, 12q, 13q, 14q, 15q, 16q, 18q, 19q, 20p, 20q) showed increased Fusion/ITS+ while 4 arms (1q, 3q, 6p, 7p) showed a decrease and one arm showed minimal change (Fig 4C).

**Table 1.** Summary of overall changes at chromosome ends in hPIF1 deficient U2OS cells.

|  | End Tel mean length, kb | Fusion/ITS+ mean length, kb | Fusion/ITS+% | TFE% |
| --- | --- | --- | --- | --- |
| siLuc | 11.16 (936/1154) | 12.51 | 17% (199/1154) | 2% (19/1154) |
| siPIF1-1 | 9.80 (711/1011) | 11.26 | 29% (298/1011) | 0.2% (2/1011) |
| P values | 0.001149 | 0.57 | 0.02 | N/A |

**Notes:** End Tel: end telomere. Fusion/ITS+: a chromosome fusion with an internal telomere-like sequence (ITS+). TFE: telomere free end.

Taken together, we conclude that hPIF1 deficiency in the ALT+ cells induces elevated replication stress and DNA damage at their telomeres, leading to a decrease in mean telomere length and an increase in telomere damaged-mediated chromosome fusions. Furthermore, our data suggest that hPIF1 may regulate telomeres differently at particular chromosome arms.

### Depletion of hPIF1 in FANCM deficient ALT+ cells attenuates the replication stress response, DNA damage, ssDNA formation, and BLM recruitment

Previously, we and others demonstrated that FANCM plays a critical role in suppressing replication stress at ALT telomeres by actively disrupting TERRA R-loops(Lu et al., 2019; Pan et al., 2019; Pan et al., 2017; Silva et al., 2019). Based on our knowledge, depletion of FANCM in multiple ALT+ cells induced the most drastic increase of almost all ALT biomarkers, including C-circles. We further showed that the drastic increase of C-circle formation in the FANCM-deficient ALT+ cells is dependent on PolD1 and PolD3, suggesting that BIR is hyperactivated in those cells(Pan et al., 2019). Next, we investigated the interplay between FANCM and hPIF1 in ALT+ cells. We co-depleted hPIF1 and FANCM in U2OS and then monitored replication stress response and DNA damage in those cells. As we previously reported, depletion of FANCM induced a drastic increase of replication stress and DNA damage at telomeres as well as a robust recruitment of BLM to the telomeres (Figs 5A to 5D)(Pan et al., 2017). Interestingly, co-depletion of hPIF1 in the FANCM deficient ALT+ cells attenuated these molecular events to half in comparison to the depletion of FANCM alone, suggesting that hPIF1 promotes the recruitment of BLM to the damaged telomeres and the activation of the replication stress response in FANCM deficient ALT+ cells. Consistently, depletion of hPIF1 also reduced C-circle formation in FANCM deficient ALT+ cells by more than 50% (Figs 5E and 5F), suggesting that depletion of hPIF1 in the FANCM-deficient ALT+ cells attenuated BIR activity by half. We observed similar reduction of recruitment of BLM to telomeres and replication stress response in Saos2 cells when hPIF1 and FANCM are co-depleted (Fig S3). However, the reduction of DNA damage in Saos2 cells is not statistically significant.

**Figure 5.**
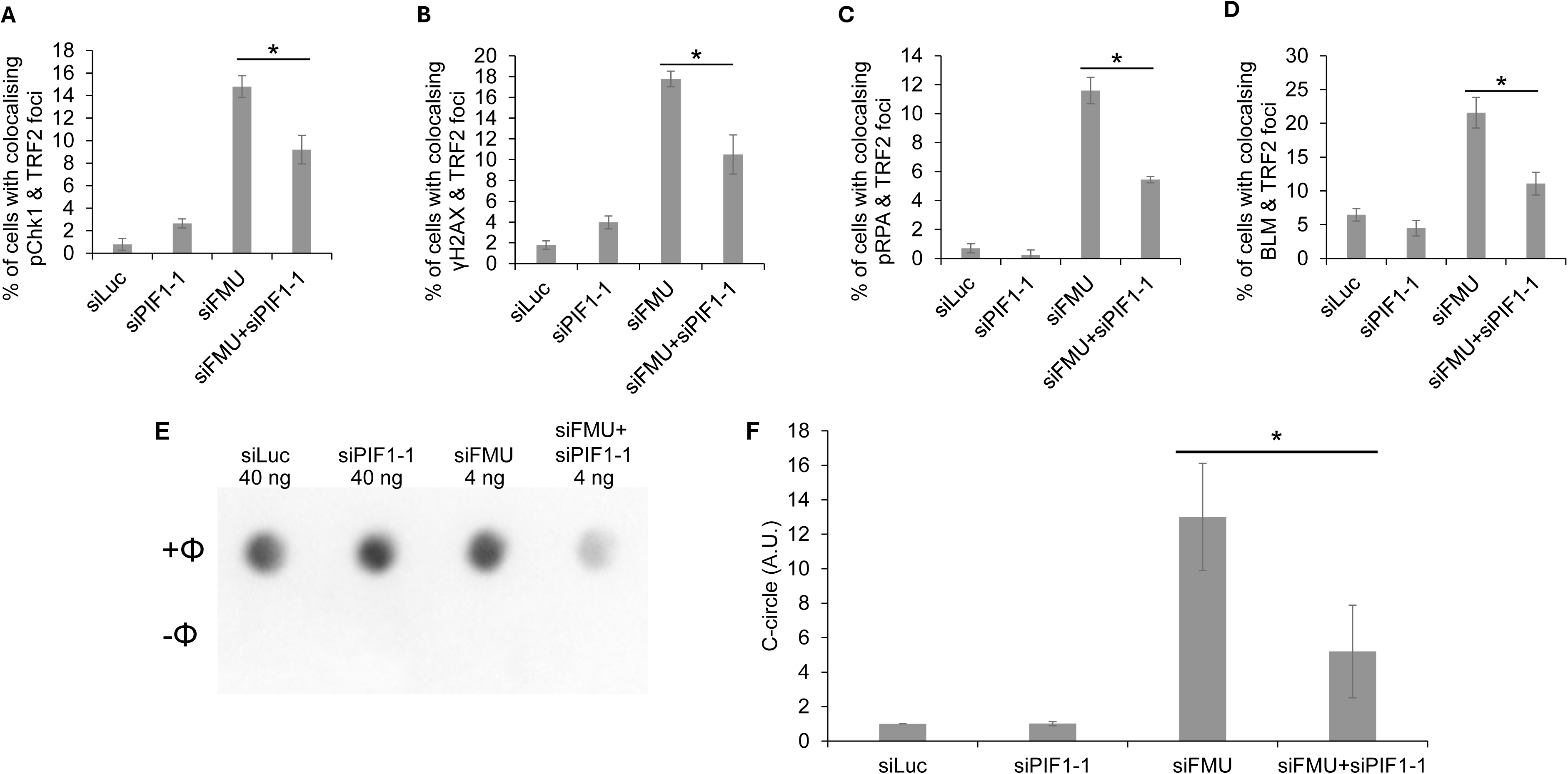
Depletion of hPIF1 attenuates replication stress response in FANCM deficient ALT cells. U2OS cells were first transfected either with a control siRNA (siLuc), a siRNA targeting hPIF1 (siPIF1-1), a siRNA targeting FANCM (siFMU), or both (siFMU+siPIF1-1). (A to D) The Cells were then stained with antibodies recognizing TRF2 and phospho-Chk1 (pChk1) (A), TRF2 and γH2AX (B), TRF2 and pRPA (C), or TRF2 and BLM (D). All nuclei were stained with DAPI. (E and F) C-circle assay in presence (+Φ) or absence (-Φ) of Phi29 DNA polymerase (Φ). All error bars are standard deviations obtained from three different experiments. Standard two-sided t test: \**p*<0.05, \*\**p*<0.01, \*\*\**p*<0.001.

In summary, our data shown here suggest that hPIF1 facilitates the activation of replication stress response, active recruitment of BLM to telomeres, and BIR in FANCM deficient ALT+ cells.

### Deficiency of hPIF1 or FANCM induces a genome-wide G-quadruplex accumulations, including telomeres

It has been well documented that PIF1 can bind and unfold G-quadruplexes (G4s) both in vitro and in vivo and this function of PIF1 is highly conserved from budding yeast to human (Malone et al., 2022). Next, we investigated whether hPIF1 and FANCM affect G4 formation in the ALT+ cells using a super resolution imaging-based G4 detection method(Fernandez et al., 2025). Inactivation of hPIF1 using either siRNA or PIF1i induced a pronounced increase of G4s at both telomeres and pan nucleus (Fig 6). Intriguingly, depletion of FANCM also induced a pronounced increase of G4s at both telomeres and pan nucleus (Fig 6). Somewhat unexpectedly, co-depletion of hPIF1 in the FANCM deficient cells attenuated formation of G4s at both telomeres and pan nucleus (Figs 6B and 6C).

**Figure 6.**
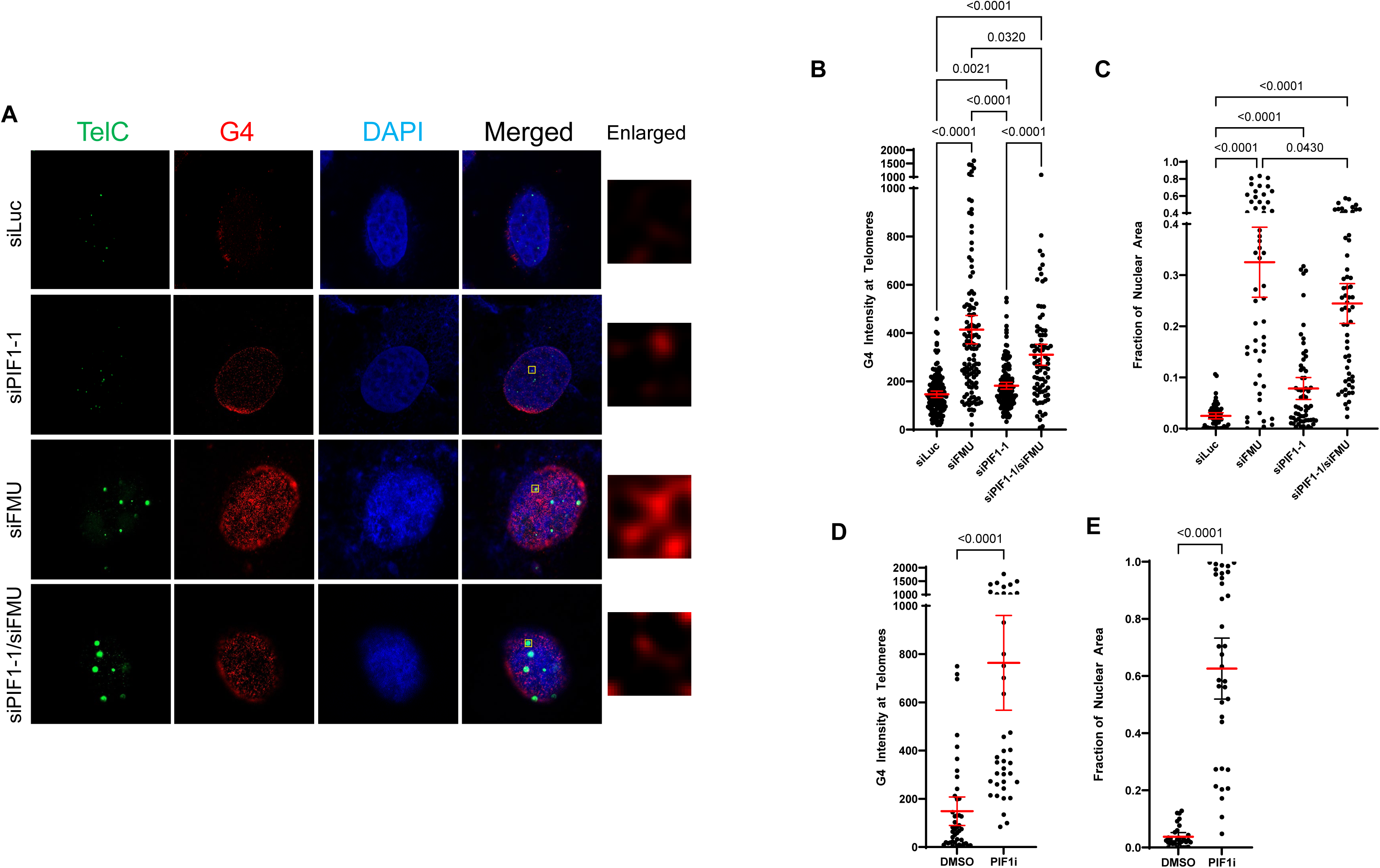
Inactivation of PIF1 or FANCM increases G4 accumulation at ALT telomeres. (A to C) U2OS cells were transfected either with a control siRNA (siLuc), a siRNA targeting hPIF1 (siPIF1-1), a siRNA targeting FANCM (siFMU), or both (siFMU+siPIF1-1). Cells were then fixed and stained with a PNA probe recognizing telomere (TelC) and an antibody recognizing the G4s. (D and E) U2OS cells were treated with either with DMSO or 300 μM PIF1i for 72 hours and then fixed and stained with a PNA probe recognizing telomere (TelC) and an antibody recognizing the G4s.

Based on our knowledge, FANCM itself does not bind and unfold G4s directly. Previously, Niedzwiedz and colleagues showed that depletion of FANCM induced a pan nuclear increase of R-loop signals in U2OS cells(Schwab et al., 2015). We and others showed that depletion of FANCM also induced a pronounced accumulation of TERRA R-loops at telomeres in U2OS(Pan et al., 2019; Silva et al., 2019). Most recently, Wu and colleagues showed that BIR can be activated to repair double-stranded DNA breaks generated from R-loops(Wu et al., 2025a). Therefore, the drastic increase of G4s in the FANCM deficient ALT+ cells is likely due to R-loop accumulation, leading to the activation of BIR which then generates excessive amount of G-rich ssDNA in many genomic regions, including telomeres. Inactivation of hPIF1 in the FANCM deficient cells attenuates BIR activity, reduces the amount of G-rich ssDNA in many genomic regions, thus decrease G4 signals at telomeres and pan nucleus.

Collectively, our data showed that hPIF1 also plays a role in unfolding G4s in vivo. In addition, our data showed the excessive ssDNA generated from the activation of BIR may promote the formation of G4s in the G-rich genomic regions, including telomeres.

### Inhibition of PIF1 in various cancer cells affect their long-term viability

Previously, Harrington and colleagues reported that PIF1 knockout mice are viable and display no visible abnormalities suggesting that PIF1 is not essential for the viability and development of mice(Snow et al., 2007). Subsequently, using the same murine model, Paquis-Flucklinger and colleagues showed that the PIF1 knockout mice develop a mitochondrial myopathy with mild respiratory chain deficiency(Bannwarth et al., 2016). Intriguingly, through analyzing clinical data sets from The Cancer Genome Atlas (TCGA), Sanders and colleagues observed that in certain cancers, overexpression of PIF1 correlates with poor patient survival(Wever et al., 2024).

To investigate whether inactivation of PIF1 affects the viability of cancer cells, we depleted hPIF1 using two different siRNA in both ALT+ (U2OS and Saos2) and TEL+ (A549, H460, and HeLa) cancer cells. Interestingly, depletion of hPIF1 affected the viability of both ALT+ and TEL+ cancer cells (Fig 7A). Consistent with these findings, when we treated these cancer cells with PIF1i, it also decreased their viability (Fig 7B). The EC_50_ of the PIF1i ranges from 60 μM to 200 μM. Collectively, our data suggests that inhibition of hPIF1 may be an effective strategy to inhibit the growth and viability of cancers while having little effect on the growth and viability of normal tissues.

**Figure 7.**
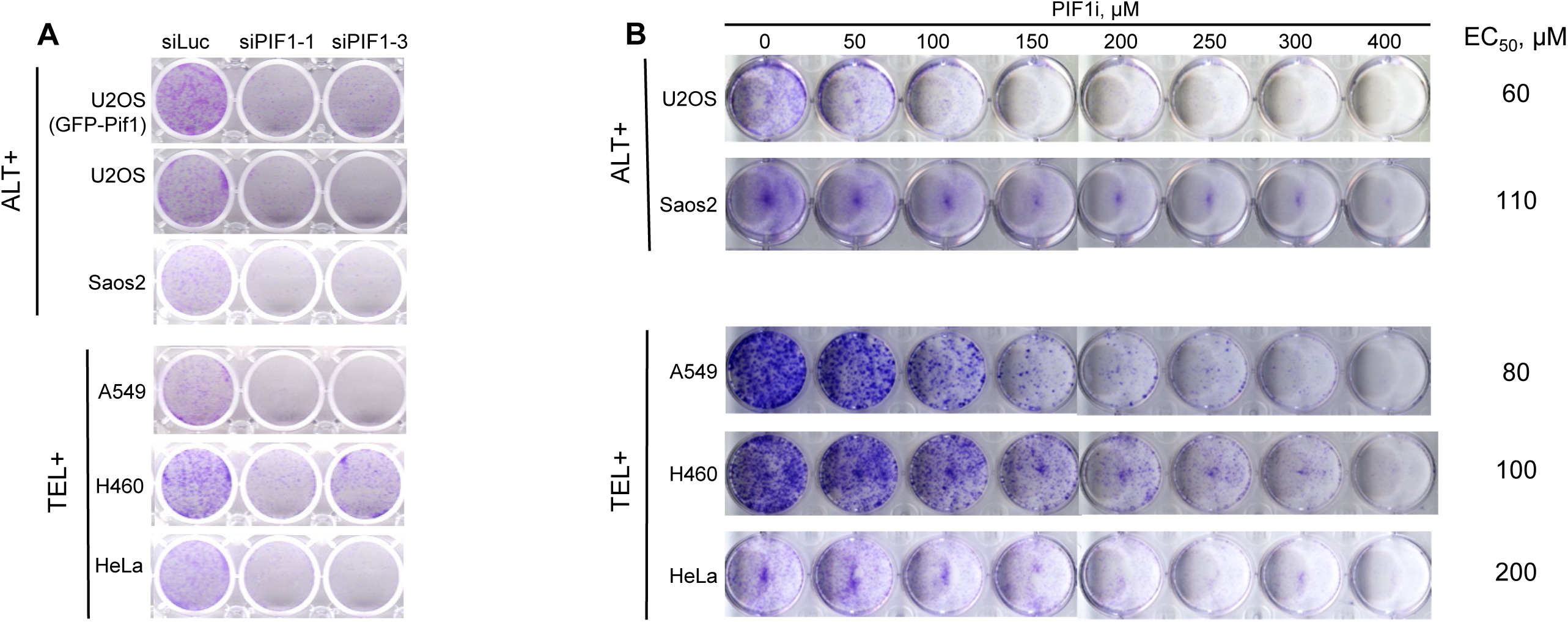
Inactivation of PIF1 affects the viability of both ALT+ and TEL+ cancers. (A) ALT+ cells (U2OS and Saos2) and TEL+ cells (A540, H460, HeLa) were transfected either with a control siRNA (siLuc), or two siRNA targeting hPIF1 (siPIF1-1 and siPIF1-3). Cells were then re-plated for Crystal Violet assay. (B) ALT+ cells (U2OS and Saos2) and TEL+ cells (A540, H460, HeLa) were treated with either DMSO or different concentrations of PIF1i and stained with crystal violet 7 to 10 days later. EC_50_ are shown on the right.

## Discussion

The budding yeast PIF1 facilitates DNA synthesis during BIR via a migrating bubble mechanism(Saini et al., 2013; Wilson et al., 2013). Using EGFP-based reporters, Wu and colleagues showed that human PIF1 (hPIF1) is also required for BIR(Li et al., 2021). However, whether and how hPIF1 is involved in the ALT pathway remains unknown. Here, we first showed that hPIF1 can be actively recruited to the damaged telomeres. We then showed that in the FANCM-proficient ALT+ cells, inactivation of hPIF1 induces a pronounced increase of replication stress and DNA damage at their telomeres and attenuates the formation of C-rich ssDNA including C-circles. In addition, we also observed a pronounced decrease of the mean telomere length and a drastic increase of telomere damage-mediated chromosome fusions. Interestingly, we found that in FANCM-deficient ALT+ cells, depletion of hPIF1 attenuates replication stress and DNA damage at telomeres as well as formation of C-circles. Furthermore, inhibition of hPIF1 or FANCM also leads to a pronounced genome-wide increase of G4 signals, including telomeres. Finally, we showed that inhibition of hPIF1 affect the long-term viability of both ALT+ and TEL+ cancer cells, suggesting that targeting hPIF1 may be a feasible strategy for cancer therapy.

### The role of hPIF1 in FANCM-proficient ALT+ cells vs in FANCM-deficient ALT+ cells

In the absence of telomerase, certain recombination-based DNA synthesis process facilitates the elongation of a shortened telomere in parental ALT+ cells (i.e., the FANCM-proficient ALT+ cells) to avoid DNA damage induced cell death. Many labs including ours showed that BIR is likely the main candidate HDR for telomere elongation when it is getting too short(O’Sullivan and Greenberg, 2025). The proposed ALT pathway to extend a shortened telomere may work as the following: in the FANCM-proficient ALT+ cell (Figs 8A and 8B), when a telomere becomes too short to form the protective T-loop, either due to the “End Replication Problem” or spontaneous telomeric damage, the exposed telomeric overhang then invades a longer telomere and uses it as the template for DNA synthesis via BIR (Scenario #1). Alternatively, there are abundant ECTRs present in ALT+ cells, including C-circles, G-circles, and a variety of linear ECTRs, which can also be used as templates to elongate the shortened telomere (Scenario #2). For example, using SMTA-OM technology, the longest linear ECTR that we can detect is around 53 kb(Raseley et al., 2023). Lastly, if linear ECTRs contain microhomology to the shortened/damaged telomere, they may be directly ligated to the telomere via the microhomology-mediated end joining (MMEJ) pathway (Scenario #3)(Singh et al., 2024). Considering the findings presented here, we propose that hPIF1 plays multiple roles at ALT telomeres. First, hPIF1 may help to actively unfold the G4s at ALT telomeres, which are formed either spontaneously (Fig 8A) or due to the formation of TERRA R-loops (Fig 8C). Secondly, hPIF1 may facilitate DNA synthesis via BIR in Scenario #1 and Scenario #2 (Figs 8B and 8C).

**Figure 8.**
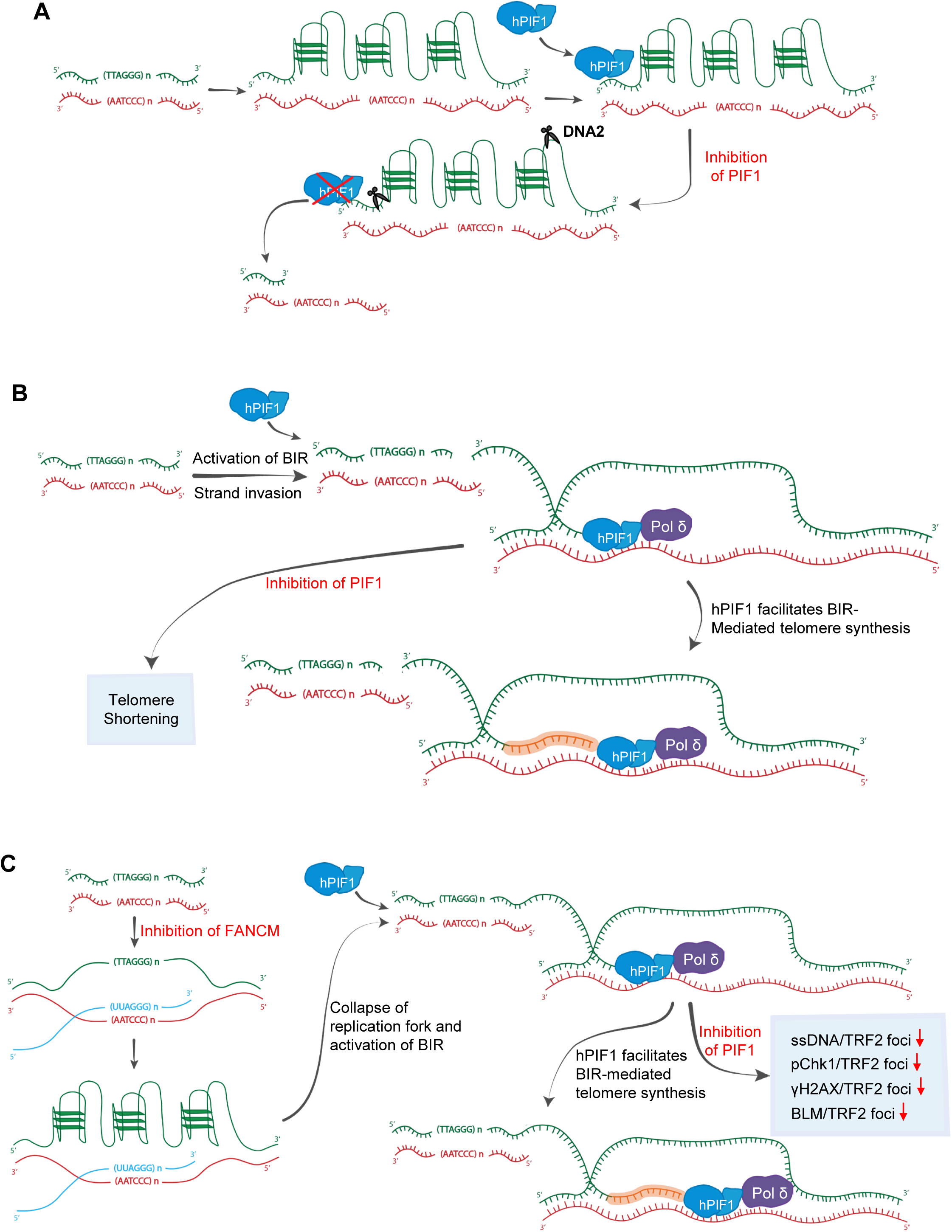
A schematic diagram of the proposed functions of hPIF1 in FANCM-proficient ALT cells and FANCM-deficient ALT cells. (A) In the FANCM-proficient ALT cells, hPIF1 helps to unfold G4s at the telomeres. Deficiency of hPIF1 leads to G4 accumulation, which then hinders DNA replication at telomeres and induce replication stress. The accumulated G4s can be cut by DNA2 or other nucleases and produces damaged telomeres. (B) In the FANCM-proficient ALT cells, hPIF1 also facilitates the activation of BIR and elongates telomeres. (C) In the FANCM-deficient ALT cells, TERRA R-loops accumulate at telomeres, which in turn promote the formation of G4s on the displaced G-rich strand. Together, the accumulated TERRA R-loops and G4s hinder DNA replication at telomeres, induce replication stress, and activate BIR, leading to the generation of excessive amount of ssDNA and further amplification of replication stress response.

In the FANCM-deficient ALT+ cells, i.e. the siFANCM transfected cells (Fig 8C), we and others showed that accumulation of TERRA R-loops at telomeres induces a drastic increase of replication stress and DNA damage (Lu et al., 2019; Pan et al., 2019; Pan et al., 2017; Silva et al., 2019). Additionally, there is also a drastic increase in C-circle formation, ranging from 20-fold to 100-fold increase depending on the detection methods. Though the mechanism of the massive increase of C-circle formation remains a mystery, we and others have shown that many DNA damage response factors and HDR factors are involved, including the BIR proteins (PolD1 and PolD3)(Lu et al., 2019; Pan et al., 2019; Silva et al., 2019). Here, we showed that hPIF1 also contributes to the generation of C-circles in the FANCM-deficient ALT+ cells (Figs 5E and 5F).

Collectively, these data strongly indicate that C-circle and other forms of ECTRs are the by-products of BIR in both the FANCM-proficient ALT+ cells and the FANCM-deficient ALT+ cells. We thus envision that the following happens in the FANCM-deficient ALT+ cells (Fig 8C): BIR is hyper-activated in the FANCM-deficient ALT+ cells due to the accumulation of TERRA R-loop-induced telomeric damage, which then generates excessive amounts of ssDNA, leading to further activation of replication stress response and DNA damage, and increase of G4 formation (Fig 6). Inhibition of hPIF1 in the FANCM-deficient ALT+ cells attenuates the activity of BIR and reduces the amount of ssDNA, thus decreases the replication stress response, DNA damage, and G4 formation (Figs 5 and 6).

### The interplays between the formation of R-loops and G4s

It has been shown that both in vitro and in vivo, hPIF1 can directly bind and unfold G4s (Jimeno et al., 2018; Sanders, 2010). Using an *in cellulo* chemically labeled pyridostatin, a well-established G4 ligand, Jackson and colleagues showed that GFP-hPIF1 colocalizes with significant numbers of pyridostatin foci(Rodriguez et al., 2012). Consistent with the previous findings, as seen in Fig 6, we observed that both siPIF1 and PIF1i induced a pronounced genome-wide increase of G4s in U2OS cells. Intriguingly, we also observed a pronounced genome-wide increase of G4s in the FANCM depleted U2OS cells. Counter intuitively, co-depletion of hPIF1 in the FANCM-deficient cells attenuated this increase. Based on our knowledge, FANCM doesn’t bind and unfold G4s directly. A previous study showed that depletion of FANCM induced a pronounced genome-wide increase of R-loops in U2OS cells(Schwab et al., 2015). We and others showed that depletion of FANCM in U2OS cells induced a robust increase of TERRA R-loops(Pan et al., 2019; Silva et al., 2019). Taken together, these data suggest that when R-loops are formed, they can promote the formation of G4s through the increased half-life of the displaced G-rich DNA strand (Fig 8C). Similarly, when BIR is hyper-active, it produces excessive amount of ssDNA and thus facilitates the formation of G4s in genomic regions that are enriched in guanines. Both accumulated R-loops and the hyper-activated BIR are likely to contribute to the genome-wide increase of G4s seen in the FANCM depleted U2OS cells (Fig 6).

### Targeting hPIF1 as a novel strategy for cancer therapy

Utilizing gene expression and clinical outcome data from The Cancer Genome Atlas (TCGA) datasets, Sanders and colleagues recently showed that there is a differential expression of hPIF1 mRNA in certain tumors in comparison to their corresponding normal tissues(Wever et al., 2024). Most importantly, they also found that there is a strong correlation between higher expression of hPIF1 and poor overall patient survival in certain cancers. Here we presented experimental evidence that hPIF1 is a potential drug target for cancer therapy. Inactivation of hPIF1 using both siRNA and a small molecule PIF1 inhibitor (PIF1i) decreased the long-term viability of both ALT+ and TEL+ cancer cells (Fig 7). Since the PIF1 knockout mice are viable and show no overt developmental defects, except mild mitochondrial myopathy(Bannwarth et al., 2016; Snow et al., 2007), an on-target PIF1 inhibitor will likely have minimal side effects when used in treating various cancers in the clinic.

## Notes

### Competing Interest Statement

The authors have declared no competing interest.

